# Wide-field Fluorescence Lifetime Imaging Microscopy with a High-Speed Mega-pixel SPAD Camera

**DOI:** 10.1101/2020.06.07.138685

**Authors:** V. Zickus, M.-L. Wu, K. Morimoto, V. Kapitany, A. Fatima, A. Turpin, R. Insall, J. Whitelaw, L. Machesky, C. Bruschini, D. Faccio, E. Charbon

## Abstract

Fluorescence lifetime imaging microscopy (FLIM) is a key technology that provides direct insight into cell metabolism, cell dynamics and protein activity. However, determining the lifetimes of different fluorescent proteins requires the detection of a relatively large number of photons, hence slowing down total acquisition times. Moreover, there are many cases, for example in studies of cell collectives, where wide-field imaging is desired. We report scan-less wide-field FLIM based on a 0.5 Megapixel resolution, time-gated Single Photon Avalanche Diode (SPAD) camera, with acquisition rates up to 1 Hz. Fluorescence lifetime estimation is performed via a pre-trained artificial neural network with 1000-fold improvement in processing times compared to standard least squares fitting techniques. We utilised our system to image HT1080 – human fibrosarcoma cell line as well as Convallaria. The results show promise for real-time FLIM and a viable route towards multi-megapixel fluorescence lifetime images, with a proof-of-principle mosaic image shown with 3.6 megapixels.

## Introduction

Fluorescence lifetime imaging, unlike conventional fluorescence imaging techniques, measures the temporal properties of a fluorophore – its fluorescence lifetime [1–3]. Fluorescence decay can be affected by the environment of the fluorophore such as concentration of oxygen, pH, or protein-protein interactions, among many others [3–5]. Hence, extracted lifetimes can reveal contrast across the sample, which would be otherwise unseen from fluorescence intensity measurements only. Fluorescence lifetime imaging microscopy (FLIM) is widely utilised in biological sciences [6, 7]. For instance, in cancer research FLIM has been used for cancer cell detection [8–11], anti-cancer or chemotherapy drug delivery [12, 13], and anti-cancer drug efficacy studies [14, 15]. In addition to this, in recent years FLIM has started to play a role in clinical diagnostics [16–18]. However, wide adoption of FLIM in clinical settings is still lacking, partially due to limited imaging speed and/or field of view (FOV) of available FLIM systems [18]. The challenges arise from the fact that nominal lifetimes of endogenous fluorophores and fluorescent proteins lie in the range of 0.1-7 ns (see Table 1 in [6]). Since there are a number of possible quenching interactions [3, 19, 20] that decrease lifetimes even further, detectors with sub-nanosecond temporal resolution are required for FLIM. Typical commercial systems make use of confocal microscopes with detectors suitable for point-scanning (such as photomultiplier tubes) and time-correlated single-photon counting (TCSPC) electronics that can satisfy the temporal resolution requirements [21]). However, point-scanning systems can suffer from photo-bleaching due to the high optical energy in the light pulses used in the system, and cannot provide instantaneous full FOV information, which becomes important when imaging dynamic scenes or *in vivo* applications. Moreover, an analytical comparison of raster scanning and wide-field data acquisition for FLIM experiments shows that for the case of very dim/sparse samples, wide-field acquisition can be *N*^2^ times faster compared to raster scanning, where *N* is the number of pixels in the detector [22]. Therefore, large detectors, such as the SPAD array used in this work, can be important for imaging dim samples which are often encountered in biologically relevant experiments.

**TABLE I.** Mean and standard deviation of extracted lifetime values of data shown in Fig. 2. The LSQ and ANN lifetime retrieval methods provide similar, compatible results.

| Data | LSQ (ns) | ANN (ns) |
| --- | --- | --- |
| HPC, Fig. 2 **a-c** | $0.90 \pm 0.18$ | $1.08 \pm 0.18$ |
| LPC, Fig. 2 **d-f** | $1.33 \pm 0.58$ | $1.30 \pm 0.29$ |
| HT1080, Fig. 2 **g-i** | $2.53 \pm 0.28$ | $2.31 \pm 0.34$ |

Wide-field FLIM is typically realised using TCSPC in a system with microchannel plate-based gated optical intensifiers combined with a sensor capable of resolving signal position, such as a Charge-Coupled Device (CCD) camera [23, 24]. An emerging alternative to aforementioned intensifier based systems are Single Photon Avalanche Diode (SPAD) arrays manufactured with complementary-metal-oxide semiconductor (CMOS) technology [25], which can operate with TCSPC or time-gated acquisition mode. The main advantages of SPAD arrays over conventional CCD/CMOS cameras are the picosecond temporal resolution, and single-photon sensitivity [26], which make them ideal for a broad range of applications in the area of ultrafast time-resolved imaging [27]. Until recently, SPAD arrays had a relatively limited number of ‘active areas’ (or ‘pixels’) due to the physical constraints imposed by the need to fit complex timing electronics for each individual pixel on the same chip. Nonetheless, recent technological advances have led to SPAD sensors with formats comparable to intensified CCDs. Prior to the development of the SPAD array used in this work[28], a 512×512 SPAD array (*SwissSPAD2* [29]) was the largest SPAD array available (recent developments in SPAD detectors are reviewed in [30]).

In this work, we demonstrate 0.5 megapixel (500×1024 pixels) wide-field FLIM microscopy with a SPAD array, while protein fluorescent lifetimes are extracted directly from the data via a bespoke artificial neural network (ANN). We illustrate the applicability of the 0.5 megapixel SPAD array for FLIM imaging of Convallaria samples at 1 Hz acquisition rate, and also in experiments with samples relevant in cancer research, such as the human fibrosarcoma (HT1080) cell line. Finally, we show the potential to extend this approach to ultra-wide fields-of-view by acquiring 8 tiles with the SPAD array, thus providing a 3.6 megapixel lifetime image of Convallaria.

### 0.5 megapixel wide-field SPAD array

The SPAD array is described in detail in Ref. [28] and in the Methods. The camera operates in gated mode, i.e. the sensor is sensitive to incoming photons for a fixed duration gate of *∼* 3.8 ns that has, to good approximation, a super-Gaussian profile (see Methods). Figure 1 **a** schematically explains how the 3.8 ns gate is scanned in steps that can be as small as 36 ps. At each step, a binary image (frame) of the spatially resolved photon counts is recorded. Stacking together all of these frames therefore provides a temporally-resolved spatial image of the sample where the data along the temporal axis is the convolution of the lifetime response with the camera temporal gate. The samples are imaged on to the camera with 0.33 µm/pixel spatial sampling for our all data, with the exception of Fig. 2 **d**-**f** that has 0.47 µm/pixel spatial sampling with (see Methods).

**FIG. 1.**
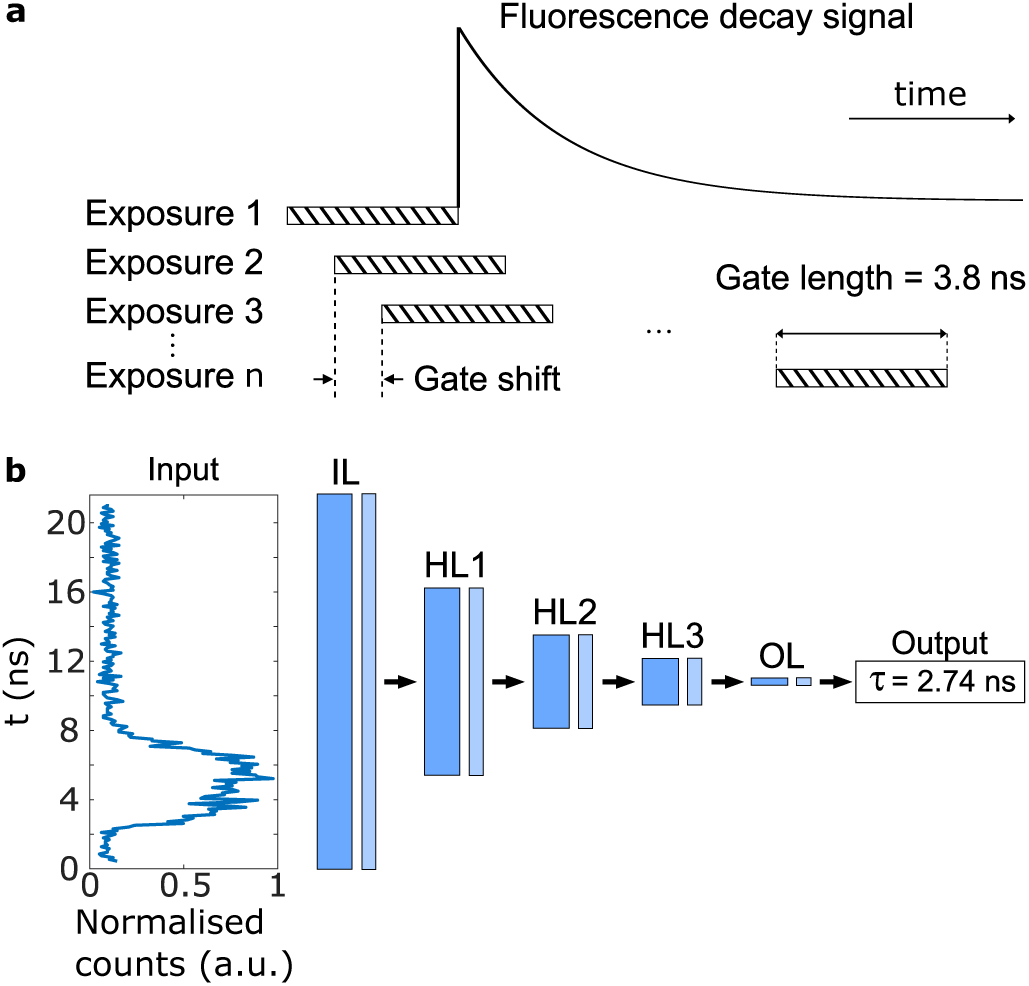
Principle of time-gated acquisition and the machine learning model. **a** Fluorescence decay is sampled with a number of gates, each shifted by a minimum of 36 ps. Each exposure corresponds to a ‘time bin’, which samples a different part of the fluorescence decay signal. **b** The ANN architecture (see Methods) consists of one input layer (IL), one output layer (OL), and a series of hidden layers (HL*i*, with *i* = 1, 2, 3). Each of these layers consists of a fully-connected dense layer (dark blue) followed by with rectified linear unit (ReLU) activation function (light blue). The input layer is fed with the fluorescence decay signal recorded by a single pixel of the SPAD array.

**FIG. 2.**
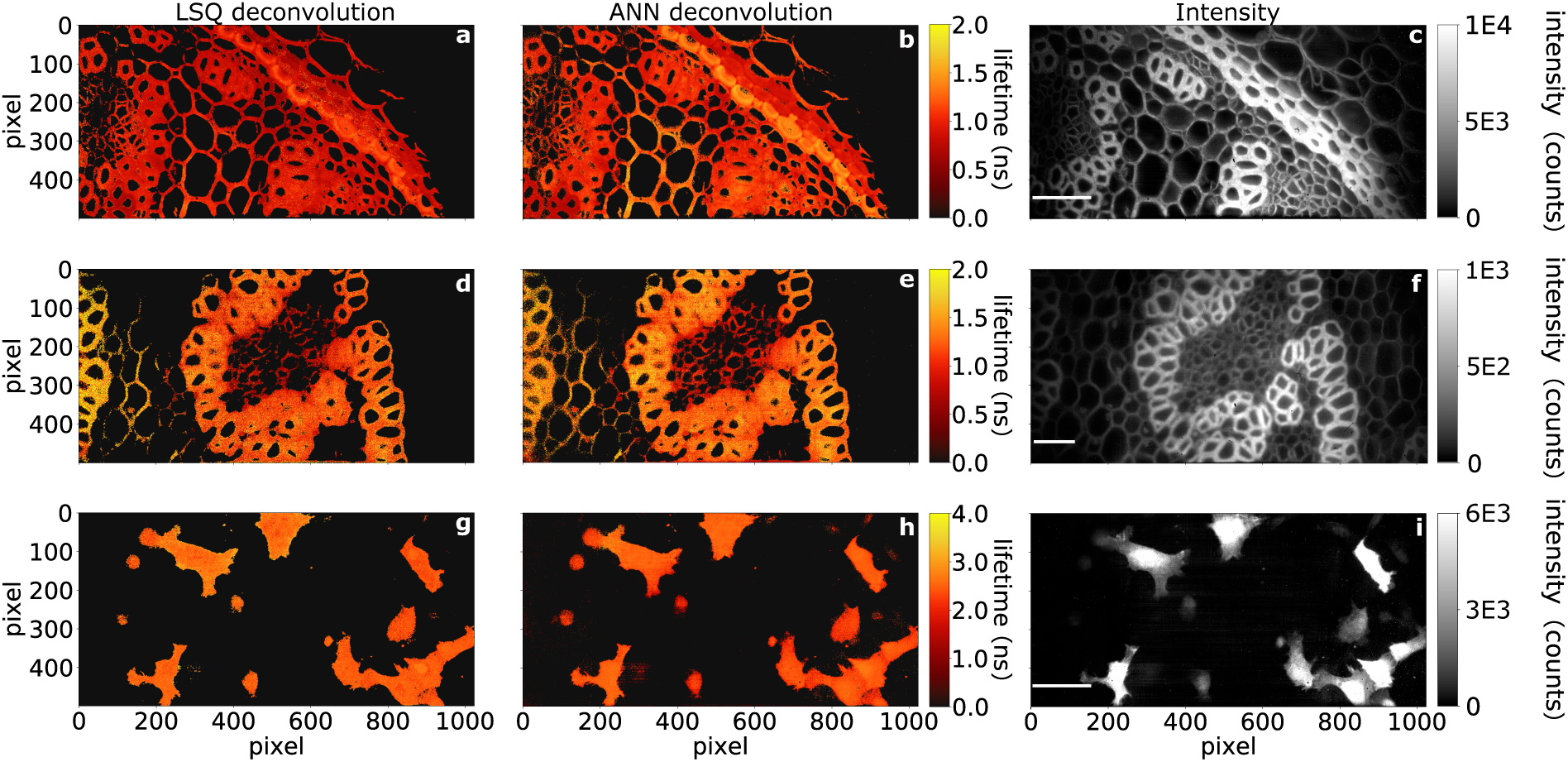
Wide-field fluorescence lifetime measurements of Convallaria and HT1080 cells. First column: least-squares (LSQ) deconvolution; second column: ANN deconvolution; third column: temporal sum of pile-up and background corrected intensity data clipped to selected intensity values to reveal dimmer structures. **a**-**c**: high photon count measurements of Convallaria (100 s acquisition). Mean lifetime measurements for LSQ (processing time, 72.3 minutes) and ANN deconvolution (processing time, 4 s) yield similar values. Spatial sampling is 0.33 µm/pixel with a 7% active area fill-factor. **d**-**f** Low photon counts measurements of Convallaria at a *total* acquisition time of 1 second. LSQ (processing time, 58 minutes) and ANN (processing time 2.7 s) deconvolution results are similar. Spatial sampling is 0.47 µm/pixel. **g**-**i** measurements of HT1080 (fibrosarcoma) cells expressing Clover [31]. As with previous data-sets with LSQ (processing time, 23.2 minutes) and ANN (processing time, 3.6 s) retrievals. Spatial sampling is 0.33 µm/pixel. Unclipped maximum intensity is above 14,000 for **c**, under 1,200 for **f**, and above 32,000 for **i**. Scale bars 50 µm.

### Lifetime retrieval

Irrespective of the imaging modality used for FLIM measurements, extracting lifetime information is not a trivial matter [32]. The measured signal, *f*(*t*), is the convolution of the impulse response function (IRF) and the fluorescence decay of the fluorophore, *g*(*t*), can be expressed as:

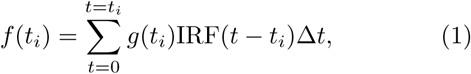

where *t*_*i*_ is the time of the *i*-th sampling of the signal. A plethora of algorithms have been developed in order to tackle the problem of retrieving fluorescence decay (reviewed in [32]), with perhaps the most common being least-squares (LSQ) deconvolution (sometimes referred to as ‘reconvolution’). In this approach a model of the fluorescence decay is convolved with the IRF and compared to the measured data using LSQ minimisation. The ‘best fit’ yields a set of parameters, including the lifetime (described in detail in Methods). However, LSQ minimisation-based lifetime estimation is typically very demanding computationally, even with the reduction of computational times provided by Graphical Processing Units (GPUs) [33]. Alternatively, fast visualisation methods such as phasor analysis have been proposed [34], and have been successfully used for time-gated SPAD array FLIM data analysis [35]. In addition to the above-mentioned numerical approaches, advances in machine learning (ML) methods [36] has enabled researchers to utilise deep learning (DL) frameworks to extract the exponential decay time and component fraction information from FLIM data rapidly and without fitting [37, 38]. Here we employ an ANN to retrieve the lifetime at each pixel of the SPAD array. The ANN layout is shown in Fig. 1 **b**: the input layer (IL) is a one-dimensional array corresponding to the time-resolved photon-count signal for a given pixel. This is connected to the output layer (OL), which provides a single value for the fluorescence lifetime decay constant *τ*, through 3 fully-connected dense layers of decreasing size. The ANN is trained on computer generated data that is created by taking a range of (mono) exponential lifetime decays (see Fig. 1 **a**) that are chosen in the range *τ* = 0.5 − 5 ns and convolved with a super-Gaussian of width that varies in the range 3.6-6 ns. The ANN was then tested on both simulated data not used in the training and also on actual experimental data. In the latter case, the retrieval was compared to the results from the LSQ deconvolution. We found that in order to retrieve precise values of *τ* from test data provided by the camera it was necessary to also include noise in the training data. The best results were obtained assuming two sources of noise: a Poisson-distributed component that is proportional to the actual signal, as expected for a photon detection process and also used in Ref. [38] and a Gaussian component whose mean (zero) and standard deviation (5 counts) were estimated from the actual data by analysing the first 10 time bins (over tens of acquisitions) in which no fluorescence signal is present. As shown in what follows, the ANN is applied to each individual pixel and can provide very similar results to a standard LSQ deconvolution albeit with a retrieval time of ∼ 8 s for a megapixel image, corresponding to 1000× gain in speed, if compared to our LSQ approach (the mean absolute difference between the two methods for the results shown in Fig. 2 was 0.14 ± 0.12 ns) and is thus a key component in rendering megapixel FLIM a real-time technique.

### Results: 0.5 megapixel wide-field FLIM

To illustrate the applicability of our SPAD array for FLIM data acquisition, we imaged a Convallaria sample with ‘high photon counts’ (HPC) at a 100 second acquisition rate (Fig. 2 **a**-**c**) and with ‘low photon counts’ (LPC) at a 1 second acquisition rate (Fig. 2 **d**-**f**). The HPC data-set was obtained by using a 108 ps gate shift, and 200 time bins. In order to achieve 1 Hz acquisition, we reduced the exposure to ≈ 33 ms per frame and we increased the gate shift to 504 ps, thus reducing the time bins from 200 to 30. Figure 2 shows that lifetime data can be retrieved for both the HPC and LPC data. However, LPC data analysis is more challenging due to the lower signal-to-noise ratio (SNR) and coarser sampling (fewer time bins) of the decay curve. In the examples shown here, the total photon count in the LPC data falls below 1,200 photons per pixel, whereas the HPC data exceeds 14,000. Nonetheless, both the LSQ and ANN methods recovered similar mean lifetime values for both HPC and LPC data. The mean lifetime and standard deviation values for the lifetime images in Figure 2 are shown in Table I.

One of the main benefits of the ANN is that it has the potential for being significantly faster than LSQ. Using a pre-trained model (training time ≈ 38 minutes on a training set of ≈ 2 million simulated decay curves), the ANN retrieval requires 2.7-4 s to process the full image. This is 500-1084 times faster than the LSQ method, which took 23.3-72.3 minutes in our tests (detailed times for each data-set are described in Fig. 2). We emphasise that we fit *each pixel* independently, and do not rely on ‘global fitting’ schemes, where data is averaged spatially and/or temporally [40].

While Convallaria is a popular sample for testing FLIM systems [41], the strong signal it yields is not necessarily representative, for example, of the signal level from transfected mamallian cells. To show a more practical example, we provide FLIM data of cancer research relevant samples: fixed HT1080 (fibrosarcoma) cells, transfected with pcDNA3-Clover [42] and expressing a protein with a single fluorescence lifetime (Fig. 2 **g**-**h**). The HT1080 cell data was acquired in HPC mode with a total 220 s acquisition time. Similarly to the Convallaria results, the ANN and LSQ results match well quantitatively (absolute difference between LSQ and ANN: 0.22 ± 0.06 ns). The large size of the sensor allows imaging multiple cells, at high detail, across a large field of view simultaneously. We note that the acquisition time could be decreased by increasing the gate shift and acquiring fewer time bins at the cost of reduced sampling of the fluorescence decay, and potentially less accurate lifetime recovery.

### Results: 3.6 megapixel wide-field FLIM

Finally, we present a 3.6 megapixel image of our Convallaria sample to showcase that very large field-of-views can be achieved almost trivially using the 0.5 megapixel SPAD array (Fig. 3). The field of view in Fig. 3 is 618×650 µm with the same spatial sampling of 0.33 µm/pixel as in previous figures. We acquired the data using a mosaic acquisition by moving the sample with ≈ 10% overlap between mosaic tiles. Crucially, our ANN method required only 36 s to recover lifetime information from this data-set. This retrieval time could be shortened by processing each pixel (or batches of pixels) in parallel.

**FIG. 3.**
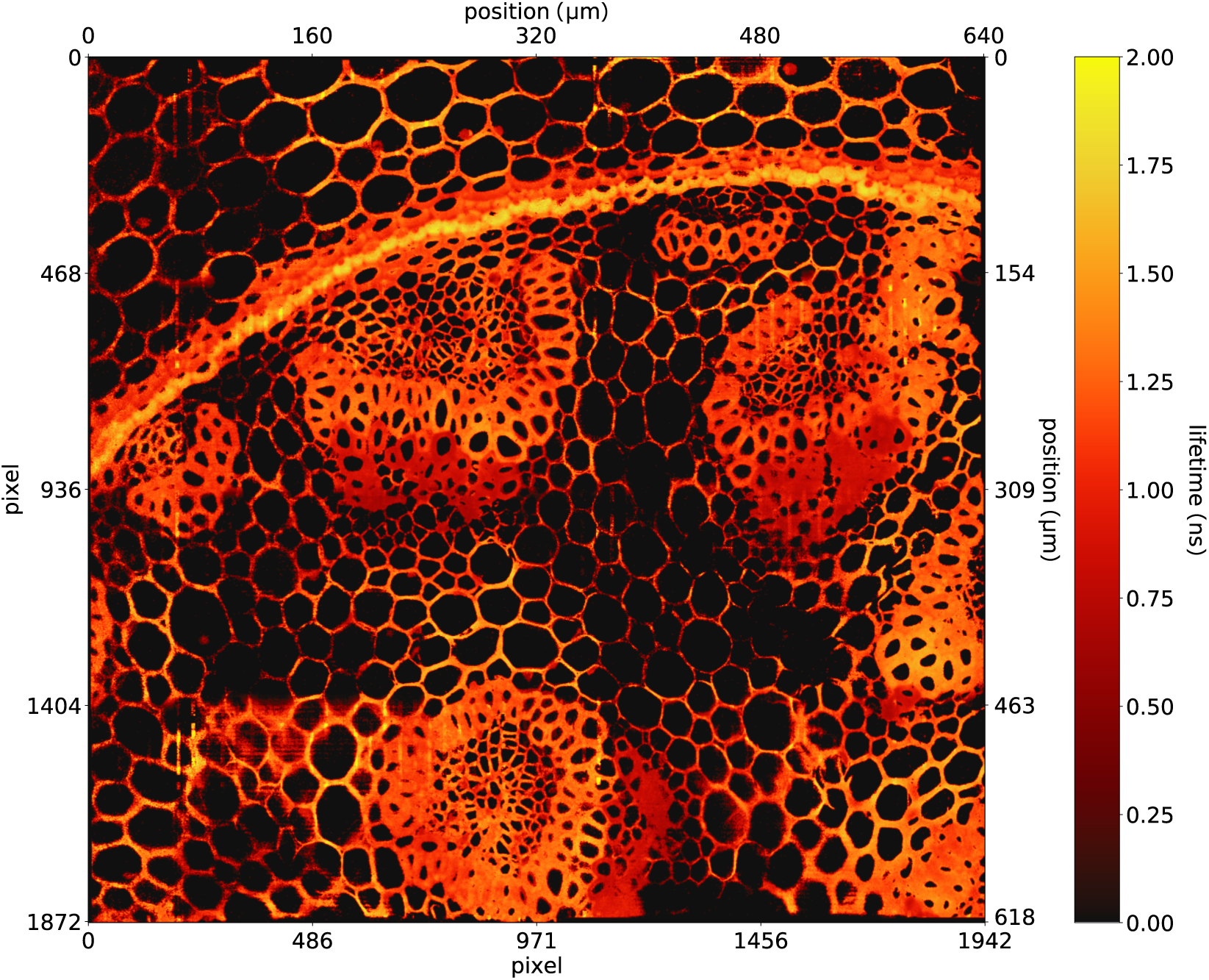
Mosaic image of 8 tiles of Convallaria sample stitched together, yielding 3.64 megapixel data (1875×1942 pixels) corresponding to a field of view ≈ 618×650 µm (or, equivalently, a sampling of 0.33 µm/pixel). The total acquisition time was approximately 16 minutes in HPC mode (that can be reduced 10-20 seconds by operating in low photon count mode) with a processing time of *≈*36 seconds using ANN deconvolution. Image stitched using *BigStitcher* [39].

### Conclusions

We have demonstrated the application of the largest-to-date time-gated SPAD array for fluorescence lifetime imaging of biologically relevant samples. By exploiting the ability to vary the gate-shift size between different exposures of the camera, we showed that wide-field FLIM at 0.5 megapixel resolution is possible at 1 Hz acquisition speed. By performing a spatial mosaic acquisition, 3.6 megapixel fluorescent lifetime images are readily available. These could be scaled to even larger fields of view and shorter acquisition times by using bespoke and rapid translation stages. In this work we retrieve mono-exponential lifetimes as this has less stringent SNR requirements compared to multi-exponential fitting [32, 43]. However, future work could of course extend this multi-exponential decays.

While fast analysis methods such as the phasor approach [34], or methods utilising advances in machine learning [37, 38], including our own ANN, are important parts of a high-speed FLIM systems, the biggest impact on imaging low-signal biologically relevant structures at sub-µm resolution at large FOV is delivered by the continuous improvement of SPAD array technology with e.g. increases also in fill factor and quantum efficiency [44].

## Acknowledgements

The authors acknowledge financial support from EPSRC (UK, grant nos. EP/T00097X/1 and EP/T002123/1), The Swiss National Science Foundation (CRSII5 177165), and Canon.

## METHODS

### Time-gated imaging

The imaging is based on a 1 megapixel SPAD image sensor[28]. The acquisition is performed by using half of the camera with 7% fill factor and 10% photon detection probability at 510 nm. To acquire the fluorescence lifetime of a sample, the camera is operated in the time-gated mode. Laser pulses are repeatedly exciting the sample; the photons emitted due to fluorescence will be detected by the camera in multiple frames with shifting gate window, resulting in an image sequence per acquisition. For each frame with a fixed gate position, 255 binary frames are summed to create an 8-bit image (9-bit images are obtained by summing two 8-bit registers). To obtain the full characteristic of a single-exponential fluorescence decay, the starting gate position is tuned to be prior to the laser excitation. The gate with average width of 3.8 ns over all pixels is then shifted finely between each frame by a fixed time. Illustration of gate-shifting is shown in Fig. 1 **a**. We note that we pre-process the data by performing background subtraction and pile-up correction, where the latter is accounted for, following equation 1 in Ref. [35] and adopting the same nomenclature:

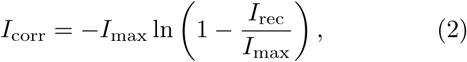

where *I*_corr_ is the pile-up corrected counts, *I*_max_ is the maximum possible photon count (depending on the bit depth e.g. 255 for 8-bit data), and *I*_rec_ is the actual recorded value at a particular pixel. The background is removed by taking the average of first few frames, before the decay signal is observed, and subtracting it from all the frames. We threshold out any pixels that have fewer than *N*_*tot*_ photon counts in total (i.e. integrated over all time gates) in order to eliminate pixels that have no significant signal. Photon counts drop towards the left hand side of the sensors due to a decay in the strength of the electronic drivers that distribute the signal controlling the time gates. We account for this non-uniform response of the SPAD array (see Fig. 5 in [28] for details) with a ‘sliding thresholding’ of the data, going from *N*_*tot*_ = 1300 (right hand edge of the image) to *N*_*tot*_ = 50 (left hand edge).

We note that despite the 7% fill factor, the small pixel pitch of 9.4 µm (corresponding to a pixel active circular area of 6.18 µm^2^) of the sensor provides sufficient sampling for microscopy at common magnification range. Figure 4 shows the Modulation Transfer Function (MTF) of our SPAD (magenta curve), indicating a 0.58 µm resolution (evaluated from the spatial frequency and half-maximum of the MTF). We also show for comparison (green curve) the MTF of a next-generation SPAD array with a higher fill factor of 42.4% and similar pixel active circular area (6.8 µm^2^) [44] and that would have more than double the resolution because of the improved pixel pitch from 9.4 µm to 4 µm.

**FIG. 4.**
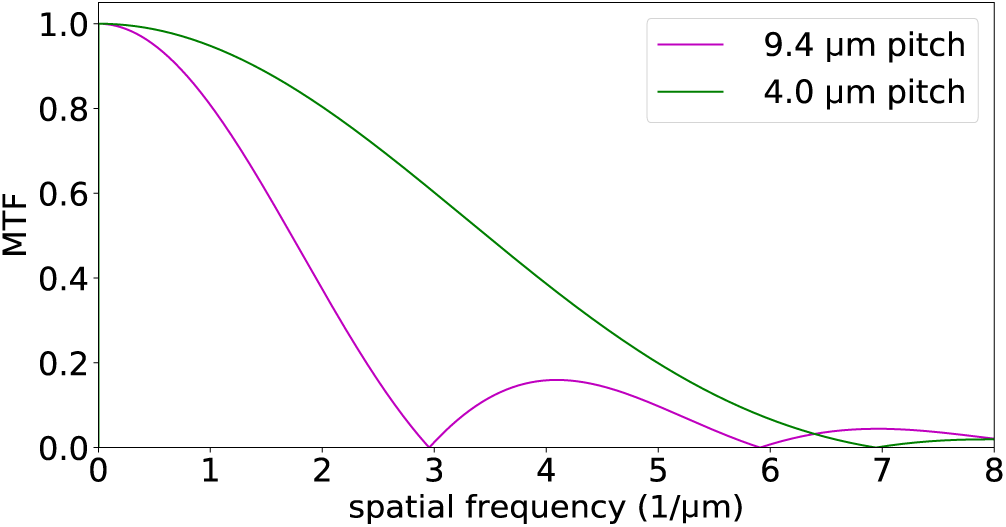
Calculated modulation transfer function (MTF) accounting for the pixel footprint, and sampling for different pixel size and pitch at an effective magnification of 27.77×, as used in our experiments. The MTF of the sensor used in this work is shown in magenta (pixel pitch 9.4 µm, fill factor 7%). Recent advances show promise for higher fill factors (42.4%) and smaller pixel pitch (4 µm) [44] thus leading to a significantly improved MTF (green curve).

**FIG. 5.**
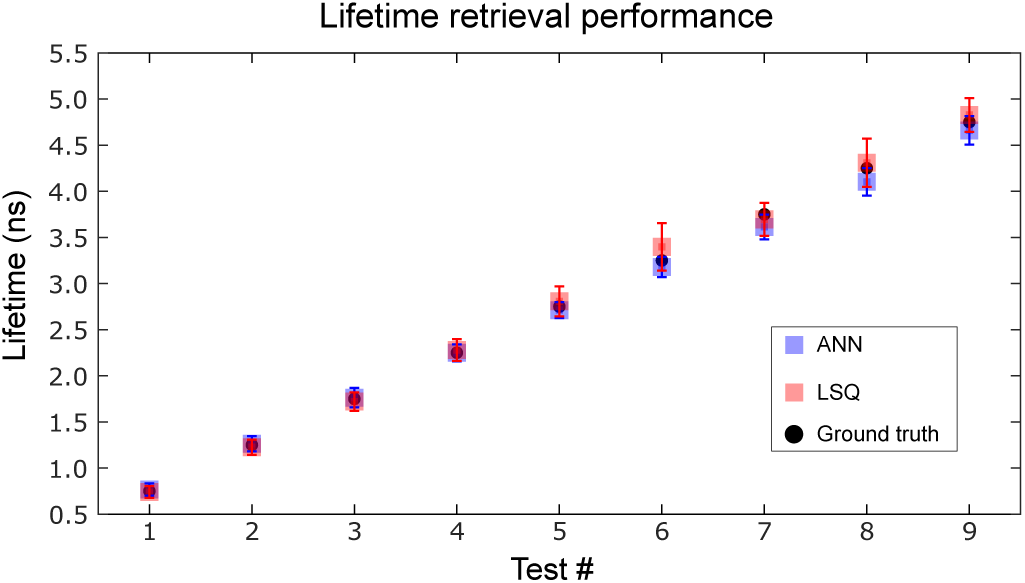
Lifetime retrieval performance of the LSQ (red squares) and ANN (blue squares) compared to the ground truth (black dots).

### Data Analysis – LSQ deconvolution

As briefly explained in the introduction in the main text, the measured data is a convolution of the impulse response function (IRF) and the underlying fluorescence decay (see Eq. 1). Technically, the IRF of a FLIM system depends on the excitation source and the detector (in our case, the gate length of each SPAD in the array). However, since the nominal pulse width of our laser pulse (*<* 47 ps) is significantly smaller than the gate length of our SPADs (average 3.8 ± 0.2 ns), the contribution to the IRF from our laser source is negligible.

We modelled the IRF as a generalised Gaussian function (or super-Gaussian) at each pixel *b* as

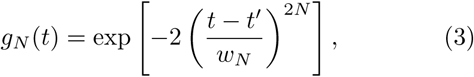

where *t* is time, *t*′ is the position of the gate, *N* the Gaussian order, and *w*_*N*_ is given by the full width at half maximum (FWHM) of the gate through the following relation:

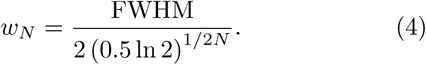

We measured that the average 10% to 90% intensity rise time for the gate (evaluated over all the pixels) is (0.55 ± 0.08) ns [28]. The order *N* of the super-Gaussian is chosen such that it matches the actual measured profiles [28]. Namely, *N* = 6 in Eq. (3) yields a rise time of approximately equal to 0.61 ns.

The fluorescence decay model used obeys the following equation:

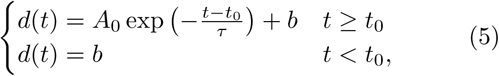

where *τ* is the decay constant (i.e. lifetime), *t*_0_ is a temporal offset, *b* is a constant that accounts for a signal offset induced by a non-zero background and *A*_0_ is an amplitude parameter corresponding to the number of photon counts. Following Eqs. (3) and (5), we model the temporal response of our detection system through the function *f*(*t*) = *g*_*k*_(*t*) ⊛ *d*(*t*), where ⊛ stands for mathematical convolution. We then apply a LSQ optimisation to the measured data that provides an estimate of [*w, t*_0_, *τ, b, A*_0_, *t*′].

### Data Analysis – artificial neural network

We use a custom ANN consisting of an input layer (IL), an output layer (OL), and three hidden layers (HL*i*, with *i* = 1, 2, 3) connecting IL with OL, as depicted in Fig. 1**b**. Each of these layers is formed by a fully-connected dense layer followed by with rectified linear unit (ReLU) activation function. The IL (with *n*_0_ = 200 nodes) is fed with a fluorescence decay signal (normalised to the range [0,1]), i.e. with a 1D vector with as many elements as the number of gate shifts (200 in our work). Then, the output of the IL is fed in cascade through the ANN, while the number of nodes of each subsequent HL*i* is decreased to: *n*_1_ = 100, *n*_2_ = 50, and *n*_3_ = 25. Finally, the OL provides an estimation of the lifetime *τ* of the fluorescence decay.

Standard machine learning techniques require training with data-sets of sufficient size and quality. To this aim, we train our ANN *only* with synthetic data, i.e. with fluorescence decay curves-lifetime pairs that are generated numerically. The training set comprised ≈ 2 million curves that are obtained by convolving fluorescence decay signals with a range of gate functions, all modelled according to Eqs. (3) and (5)). As described in the main text, we also included a noise model in the data, as this proved to be essential in retrieving precise lifetime values on test data. Including variability on gate width, centre position, lifetime, and lifetime curve height, allows us to account for the fact that the various SPAD pixels have slightly different underlying electronic properties [28]. After the ANN is trained and tested on unseen synthetic data (see below), it is fed with fluorescence decay curves experimentally recorded by individual pixels from the SPAD array in order to estimate the lifetime, *τ*.

Both LSQ and ANN deconvolution approaches retrieve fluorescence lifetimes one pixel at a time, and can therefore be used on data of any dimension. For the ANN approach we obtained a root mean squared error of 0.0725 on a test set of ≈ 1 million synthetic data. The ANN algorithm then takes (≈ 8.5 ± 0.5) s to predict the lifetimes of a 1024×1024 synthetic data set with 200 time bins on a Intel Core i7 10510U CPU.

Fig. 5 shows the lifetime retrieval performance of both the LSQ and the ANN methods. For this benchmark, we generated 9 sets of synthetic curves following the convolution model described above. Each set of curves had constant lifetime decay *τ* within the range [0.75 − 4.75] ns (black dots in Fig. 5) and randomly varying noise from curve to curve within the set. Red and blue squares represent the mean of the lifetime values retrieved with the LSQ and the ANN, respectively, while error bars correspond to their standard deviation. Both methods provide lifetime estimates in excellent agreement with the ground truth data.

### Imaging set-up

We acquired the data on a custom-built epi-fluorescence microscope, with an Olympus 20×0.4NA objective with an *f* = 250 mm tube lens (or Nikon 400.×75NA objective, and *f* = 100 mm tube lens for 1 Hz acquisition) air objective, and a FITC emission/excitation filter and dichroic mirror set. For the illumination source, we used HORIBA DeltaDiode laser diode model *DD-470L*, nominal peak wavelength 470 nm spectral Full Width at Half Maximum (FWHM) *<* 10 nm, nominal pulse width of *<* 47 ps, and a nominal pulse energy of 15 pJ. The repetition rate of the diode was set to 25 MHz.

### Mammalian cells, culturing conditions and transfections

HT1080 cells were maintained in Dulbecco’s modified eagle’s medium (DMEM), supplemented with 10 % foetal bovine serum (FBS), 2 mM L-glutamine and 1X PenStrep. Cells were maintained in 10 or 15 cm TC-treated plastic dishes at 37°C and 5 % CO_2_. HT1080 cells were transfected using Amaxa nucleofector Kit T, program L-005. Cells were transfected with 5 µg DNA (pcDNA3-Clover) following manufacturers guidelines and replated on 6 cm TC-treated plastic dishes overnight at 37°C, 5 % CO_2_. 35 mm glass bottom Mat-Tek dishes that were coated with laminin 10 µg ml^*−*1^ diluted in PBS and left overnight at 4°C. Cells were then collected and replated onto the dishes and incubated for 24 hours at 37°C, 5 % CO_2_. After, the dishes were washed twice with PBS before fixing with 4 % Paraformaldehyde for 10 minutes. These were then washed three times with PBS before slight drying. 10 µl Fluromount-G (without DAPI) was added on top of the cells and a 19 mm glass coverslip was added on top to seal the cells in the dish. Dishes were kept in the dark until imaging.

